# Optical Co-registration of MRI and On-scalp MEG

**DOI:** 10.1101/498113

**Authors:** Rasmus Zetter, Joonas Iivanainen, Lauri Parkkonen

## Abstract

**Objective:** To estimate the neural current distribution underlying magnetoencephalographic (MEG) signals and to link such estimates to brain anatomy, MEG data have to be co-registered with an anatomical image, typically an MR image. Optically-pumped magnetometers (OPMs) enable the construction of on-scalp MEG systems providing higher sensitivity and spatial resolution than conventional SQUID-based MEG systems. Here, we present a co-registration method that can be applied to on-scalp MEG systems, regardless of the number of channels.

**Methods:** We apply a structured-light 3D scanner to create a surface mesh of the subject’s head and the sensor array, which we fit to the MR image. To assess accuracy, we quantified the reproducibility of the surface mesh and localized current dipoles embedded in a phantom. Finally, we measured somatosensory evoked fields (SEF) to median nerve stimulation and compared their source estimates with those obtained with a SQUID-based MEG system.

**Results:** The structured-light scanner reproduced the head surface with < 1 mm error. Phantom dipoles were localized with a mean error of 2.14 mm. The difference in SEF dipole positions between OPMs and SQUIDs were 5.0, 0.9, and 1. 6 mm for N20m, P35m and P60m response peaks, respectively.

**Conclusion:** The developed co-registration is inexpensive, fast and can easily be applied to on-scalp MEG. It is also more convenient to use than traditional co-registration methods while also being more accurate.

**Significance:** We developed and validated a co-registration method that can be applied to on-scalp MEG systems. The method enables accurate source estimation with these novel MEG systems.

## I. Introduction

Magnetoencephalography (MEG) is a non-invasive functional neuroimaging method for investigating electric neuronal activity inside the living human brain [1]. MEG systems measure the magnetic field produced by neural currents in the brain using sensors positioned around the head.

So far, the sensor type employed for MEG has almost exclusively been low-T_c_ superconducting quantum interference device (SQUID). Low-T_c_ SQUIDs require a cryogenic temperature that is typically attained by immersing SQUIDs in liquid helium (*T* ≈ 4.2 K ≈ – – 269°C). The necessary thermal insulation keeps SQUIDs at least 2 cm away from the scalp in most commercial systems and makes the construction of adaptable sensor arrays extremely challenging. Since sensitivity and spatial resolution are related to the distance between the sources and the sensors, the need of cryogenics eventually results in a considerable loss of signal amplitude and spatial resolution [2], [3].

New sensor technologies with sensitivity high enough for MEG have emerged recently; optically-pumped magnetometers (OPMs) [4], [5] and high-T_c_ SQUIDs [6] hold promise as alternatives to low-T_c_ SQUIDs. These new sensor types do not require the same degree of thermal insulation as low-T_c_ SQUIDs and can thus be placed almost directly on the scalp, considerably boosting both the sensitivity to neural sources as well as spatial resolution.

In order to determine the origins of neuromagnetic signals measured with MEG, one needs to incorporate anatomical information, typically obtained from structural MRI. To be able to combine data from the two imaging modalities, their coordinate systems need to be aligned, i.e., co-registered. Accurate co-registration is particularly important in on-scalp MEG [7] due to its high spatial resolution [2], [3].

The current standard co-registration method in SQUID-based MEG relies on the combination of head position indicator (HPI) coils attached to the participant’s head and a pen-like electromagnetic 3D digitizer. Prior to MEG measurements, the positions of the HPI coils as well as a set of anatomical landmarks on the head are digitized. To localize the HPI coils with respect to the MEG sensors, known currents are driven into the coils either sequentially or at different frequencies prior to or continuously during MEG measurements and a magnetic dipole model representing each coil is fitted to the acquired MEG sensor signals.

The accuracy of the co-registration can be improved by digitizing not only the landmarks but a larger set of point on the head surface. Due to the need to manually digitize each point, their number is limited. For the same reason, the number of HPI coils is typically no more than 5. However, with optical scanning methods, one can obtain several orders of magnitude more points in less time than used with current methods. Additionally, the accurate localization of the HPI coils requires a MEG sensor array with extensive coverage and a large number of channels. Thus, using HPI coils with the current, early-stage on-scalp MEG systems with only a few channels is not feasible.

Here, we describe a co-registration method that employs a commercial, consumer-grade structured-light scanner and is suitable for an on-scalp MEG system with a partially rigid sensor array. We validate the co-registration method both in terms of reproducibility and accuracy, using phantom measurements as well as a human experiment.

## II. Materials and Methods

### A. Structured-light scanner

We applied a consumer-grade structured-light scanner (Occipital Inc., San Francisco, CA, USA) to digitize the subject head surface as well as the MEG sensor helmet. The structured-light scanner functions by projecting a pattern of infrared light onto an object, which is then detected by a camera at a known distance from the projector. The three-dimensional shape of the scanned object can then be determined based on the apparent distortion of the pattern as seen by the camera. The scanner captures both color and depth data at a frame rate of 30 Hz, with each frame being co-registered to the previous one in real time. The scanner is connected to a tablet (iPad; Apple Inc., Cupertino, CA, USA), which can both function as an operator display and compute the digitized surface mesh in real time. At a typical working distance of 50 cm, the scanner has a vendor-specified point accuracy of 0.8 mm [8].

### B. On-scalp MEG system

We applied the structured-light scanner to an on-scalp MEG system comprising nine QuSpin ZF-OPM sensors (QuSpin Inc., Louisville, CO, USA) in a partially rigid array, where each sensor measures the magnetic field component approximately normal to the head surface. For a detailed description of the system, including data acquisition electronics, see [9]. The sensors are mounted in a 3D-printed helmet with geometry identical to that of the Elekta Neuromag® Vectorview and TRIUX (MEGIN / Elekta Oy, Helsinki, Finland) 306-channel SQUID-based MEG systems. Individual OPMs are placed into sockets, whose positions and orientations correspond to those of the above SQUID systems, and inserted until touching the head of the subject. The insertion depth is manually measured for each sensor.

For the human measurements, the helmet was attached to the subject chair, and two dummy sensors were used to fix the subject’s head to the sensor helmet in order to minimize movement (Fig. 1, top panel).

**Fig. 1:**
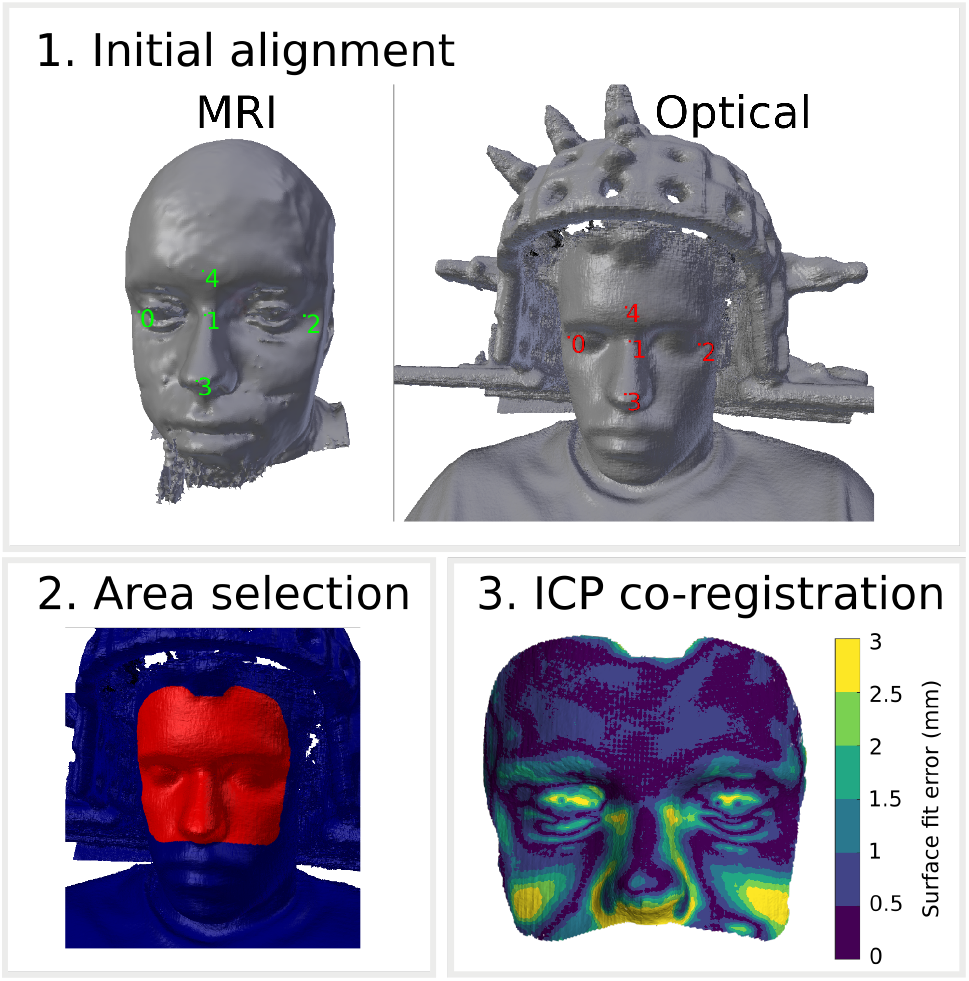
Optical co-registration procedure: 1. Initial mesh alignment with manually selected fiducial points on the MR (left) and structured-light scan (right) meshes. 2. Selection of the co-registration area. 3. Automatic ICP-based co-registration and visualization of the result.

### C. SQUID-based MEG system

For validation and comparison purposes, MEG was also recorded using a 306-channel SQUID-based Elekta Neuromag® VectorView MEG system. Data were recorded using a sampling frequency of 1 kHz. For the analyses in this work, only the 102 magnetometers in the system were used as to provide a measurement directly comparable to that of the OPM-based MEG system.

### D. MEG-MRI co-registration process

After positioning the subject in the OPM-MEG system, an optical scan is performed by moving the structured-light scanner around the subject at a distance of approximately 50 cm. The digitized surface is visualized in real time, enabling the operator to perform quick corrections and fill in any gaps in scan coverage by going over specific areas again. The scan takes approximately 30 s to perform, depending on the desired coverage. While the real-time constructed mesh functions as a good reference, it has limited resolution. For increased accuracy and resolution, we apply an offline mesh reconstruction algorithm found in the Skanect software package (Occipital Inc., San Francisco, CA, USA) to compute the final surface mesh.

Using the *mkheadsurf* tool included in the FreeSurfer software package [10]–[12], a corresponding scalp surface mesh is extracted from the structural MR image.

The co-registration algorithm was implemented as a plug-in to Blender (https://blender.org), an open-source 3D creation software suite. The co-registration process is initialized by a rough alignment of the optical scan mesh and the MR scalp mesh by manually selecting and aligning a small number of fiducial points on both meshes (Fig. 1, top panel).

Since the optical scan includes both the subject’s head and the MEG helmet with sensors, the areas (vertices of the meshes) used for co-registration with the MR image must be restricted. This is done manually by “painting” over the desired parts of the mesh with the computer mouse (Fig. 1, bottom-left panel). Any parts of the optical scan can be explicitly excluded in the same manner. Being able to quickly constrain the areas used for co-registration makes it easy to exclude areas with visible artifacts. For co-registration of the MR scalp mesh and optical scan mesh, only the upper face and forehead areas are used. Previous work has shown that using face data for high-density meshes is sufficient for accurate co-registration of EEG and MEG [13]–[15].

After the initial alignment and vertex selection, a variant of the iterative closest point (ICP) algorithm with point-to-point minimization is run for up to 100 iterations to automatically co-register the meshes. If the mesh translation is less than 0.1 mm in an iteration, the algorithm is stopped (typically within 10 iterations, Fig. 1, bottom-right panel).

After co-registering the optical scan and the MR image, the same procedure is repeated to align the known geometry of the MEG helmet to the optical scan (now in MR coordinates). The coordinates and orientations of the sensor slots of the helmet are then retrieved from the helmet model, and the manually-measured insertion depths are taken into account to get the actual sensor positions.

## III. Experiments

### A. Phantom experiment: Reproducibility

To attain a baseline scanner performance estimate, we scanned a head-shaped polystyrene phantom five times, with each scan taking approximately 45s. These five repetitions were thereafter co-registered using the process described above. After co-registration, we used the initial scan as a baseline and for each vertex we computed the distance to the closest vertex in the subsequent scans.

### B. Phantom experiment: Dipole localization using OPMs

To validate the co-registration methodology in a controlled manner, we applied a calibrated dry phantom (MEGIN / Elekta Oy, Helsinki, Finland) containing 32 sources at known positions within a head-sized hemisphere. Each source is a current loop that comprises a 5-mm long tangential segment producing a field of a current dipole and two radial segments that generate a field corresponding to the volume-current field associated with that current dipole. Thus, the total field is that of a current dipole in spherical conducting medium.

Eight of these sources were sequentially energized with 2 cycles of a 20-Hz sinusoid with a peak amplitude of 500 nAm. This pulse was repeated 100 times while the produced magnetic field was measured using the OPM-MEG system. The sensors were positioned as to cover both negative and positive field maxima. This measurement procedure was repeated five times with slightly different positions of the phantom with respect to the sensors. For each repetition, the phantom and sensor helmet were co-registered using the structured-light scanner.

### C. Human experiment: SEF measurements using OPMs and SQUIDs

We recorded somatosensory evoked fields (SEFs) from a single subject (male, 28 years) using both the SQUID-based 306-channel and a 9-channel OPM-based MEG systems. Somatosensory responses were produced by transcutaneous electric median nerve stimulation delivered to the left median nerve at the wrist using 7.5-mA 200-*μ*s current pulses. 200 trials were recorded with an inter-trial interval of 2.5 s with jitter uniformly distributed to ± 0.2 s.

For the SQUID measurements, co-registration was performed prior to MEG recording using HPI coils and a Polhemus®Isotrak electromagnetic digitizer. For the OPM-based system, the structured-light scanning co-registration presented in this work was applied. The OPMs were placed above the somatomotor areas of the hemisphere contralateral to the stimulation, such that they would measure both the positive and negative field maxima while also providing good spatial resolution (Fig. 4).

A T1-weighted structural MR-image was used in source modeling. The FreeSurfer software package [10], [12], [16] was used for pre-processing the MRI and for segmentation of the cortical surfaces. The skull and scalp surfaces were segmented using the watershed approach [17] implemented in FreeSurfer and MNE software [18]. These surfaces were thereafter decimated to obtain three boundary element meshes (2562 vertices per mesh).

MEG data were filtered to 0.1–100 Hz in both measurements, epochs were manually inspected for artifacts, and thereafter averaged. Single dipoles were fitted to the first three response peaks (N20m, P35m and P60m). All analysis was performed using MNE-Python software [18]. Dipole fitting for the SQUID measurements was performed using only a subset of the 102 magnetometers, retaining only those channels that showed some evoked activity (see Fig. 4).

## IV. Results

### A. Phantom experiment: Reproducibility

The mean reproducibility error of the optical scanner was 0.87 mm across all repetitions and vertices (error distribution modes within 0.42–0.49 mm). As is evident from Fig. 2, the errors are larger in the area under the chin of the phantom head, which the scanner could image only at very oblique angles. This area with larger errors is also seen in the error distributions in Fig. 2 as the right-hand-side tail.

**Fig. 2:**
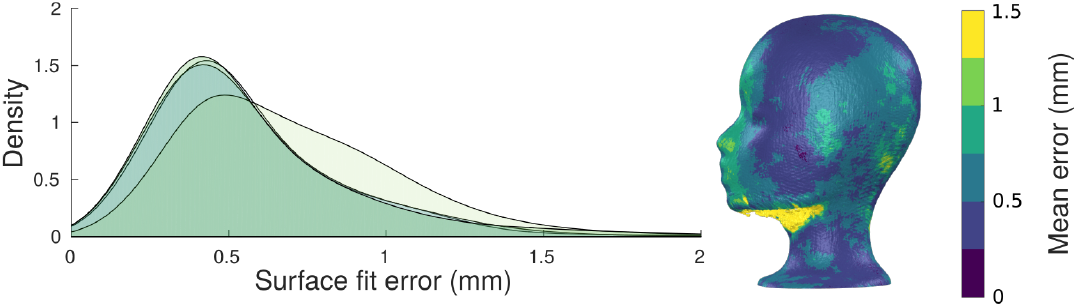
Reproducibility of the surface mesh reconstructed by the optical scanner. Distributions (left) and spatial locations (right) of errors across five scans of the same object.

### B. Phantom experiment: Dipole localization using OPMs

The accuracy of localizing the dipolar current sources in the phantom is shown in Fig. 3 (top panel). Across all dipoles, the absolute localization error was 2.14±1.07 mm (mean±SD), the orientation error was 2.63°±1.22° (mean±SD) and the amplitude error was −10.27±21.17 nAm (the nominal amplitude was 500 nAm). Goodness-of-fit varied within the range 94.88−99.92% (98.98±0.96%, mean±SD).

**Fig. 3:**
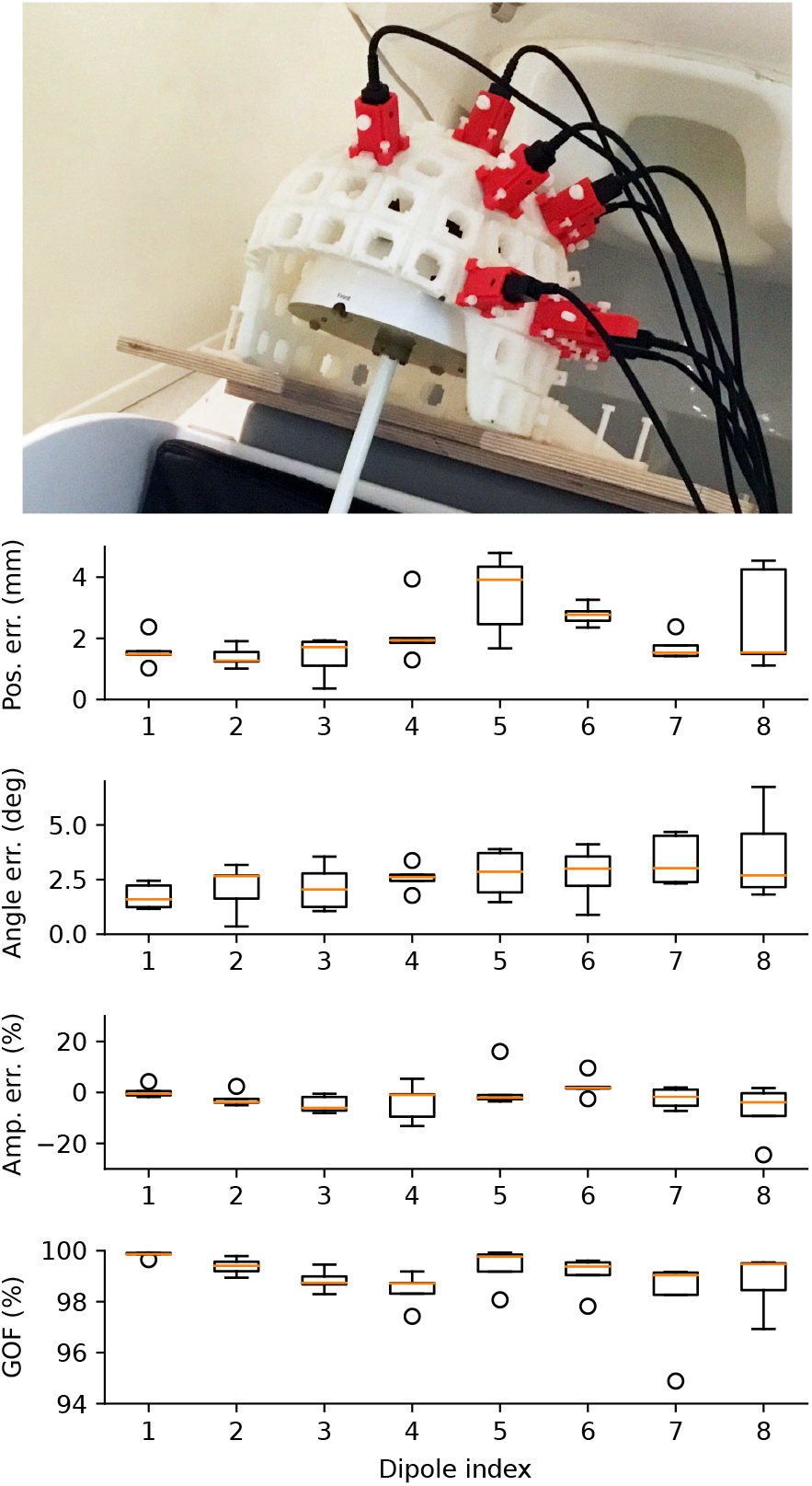
Top:Phantom measurement setup. Bottom: Phantom dipole localization; absolute position, orientation and amplitude errors as well as goodness-of-fit (GOF).

**Fig. 4:**
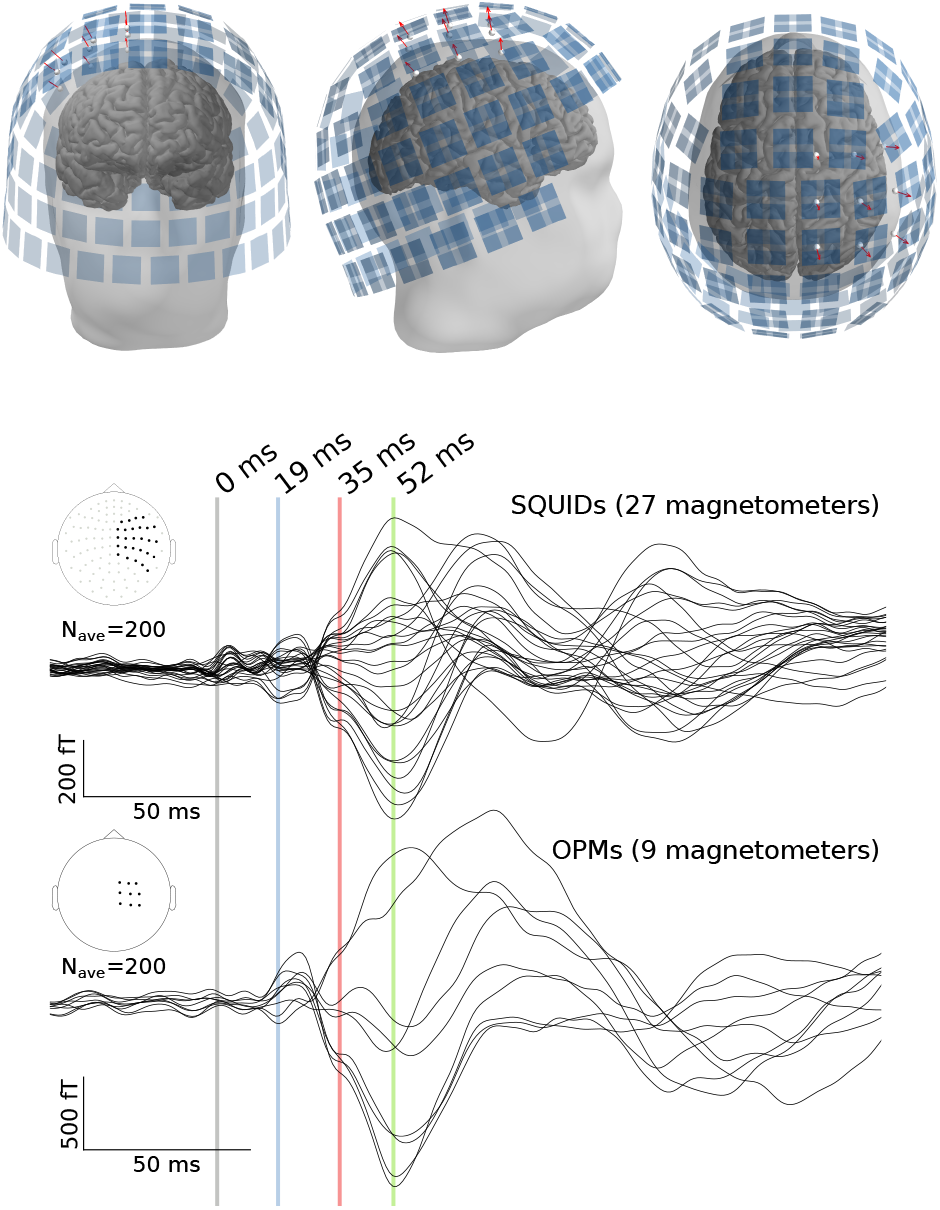
Top: Positions of the OPM (white spheres; sensitive axes as red arrows) and SQUID (blue rectangles) sensors in the somatosensory measurement. Bottom: Somatosensory responses in the OPM and SQUID (subset as shown in the array layout) channels. Equivalent current dipoles were fitted at the indicated latencies.

### C. Human experiment: SEF measurements using OPMs and SQUIDs

The matching of the optical scan and MRI meshes of the test subject is illustrated in Fig. 1 (bottom-right panel).

Equivalent current dipoles were fitted to the peaks of the three earliest responses;19 ms (N20m), 35 ms (P35m) and 52 ms (P60m) for both OPM- and SQUID-MEG measurements. Characteristics of the fitted dipoles are shown in Table I.

**TABLE I:** SEF equivalent current dipole parameters for the three earliest response peaks.

| Dipole | Time [ms] | System | Amplitude (nAm) | GOF (%) | Position (x, y, z) (mm) | ΔPos (mm) | Orientation (x, y, z) | ΔOri ($^\circ$) |
| --- | --- | --- | --- | --- | --- | --- | --- | --- |
| N20m | 19 | OPM | 10.4 | 90.3 | 43.0, -15.6, 74.8 | 5.0 | -0.35, 0.88, 0.31 | 7.7 |
|  |  | SQUID | 14.7 | 80.4 | 42.3, -11.3, 72.6 |  | -0.46, 0.81, 0.35 |  |
| P35m | 35 | OPM | 15.7 | 95.3 | 41.3, -22.6, 80.0 | 0.9 | 0.44, -0.80, -0.40 | 0.4 |
|  |  | SQUID | 15.2 | 89.6 | 41.4, -23.2, 78.7 |  | 0.44, -0.80, -0.40 |  |
| P60m | 52 | OPM | 65.9 | 99.7 | 42.8, -21.1, 73.0 | 1.6 | 0.44, -0.78, -0.44 | 6.3 |
|  |  | SQUID | 45.4 | 97.0 | 44.3, -21.0, 73.6 |  | 0.51, -0.71, -0.48 |  |

## V. Discussion

Accurate co-registration of MEG and MRI is critical for reliable source estimation as shown in several studies concerning conventional SQUID-based MEG [19], [20], EEG [21], [22] and more recently also on-scalp MEG [7]. In this work, we present a co-registration method based on a structured-light scanner that can be applied to on-scalp MEG, but could also in principle be applied to SQUID-based MEG.

The traditional co-registration procedure based on HPI coils and an electromagnetic digitizer has been challenged during the past ten years by demonstrating faster or more accurate digitization and co-registration methods (e.g. [14], [15], [19], [21], [23], [24]). With the ongoing development and commoditization of consumer-grade 3D scanners, the use of optical co-registration has become increasingly attractive.

The accuracy of the optical scanner employed in this work seems to be satisfactory, with error levels at the sub-millimeter level (Fig. 2), which is on par to what is specified for the popular Polhemus®electromagnetic digitizer [25]. However, the weakness of digitizer-based co-registration methods is their manual, operator-dependent nature; a human has to accurately move the digitizer to each reference point (without shifting the point itself). The digitization of HPI coils is especially sensitive to operator error, as only five or even fewer HPI coils are typically used. In practice, the reported accuracy of Polhemus-based co-registration ranges from 3 to 7 mm [14], [15], [24], [26].

Optical co-registration methods also have limitations, the foremost of which is the line-of-sight issue. OPM sensors are typically closely spaced on the scalp, and together with their cables they present a complex surface that is not easy to digitize accurately, with complete coverage and without artifacts. Therefore, we register the rigid sensor helmet to MRI, and simply measure the insertion depth of each sensor into the helmet instead of digitizing the location of each sensor directly.

Furthermore, when using optical co-registration, one needs to identify each sensor unambiguously. In our case, the use of a sensor helmet with discrete slots simplifies the matter; we log the assignment of sensors to the helmet slots for each measurement. In earlier EEG work, automated algorithms [13], [27], colored markers [28] or even timed LEDs [29] have successfully been used to solve this problem.

**Fig. 5:**
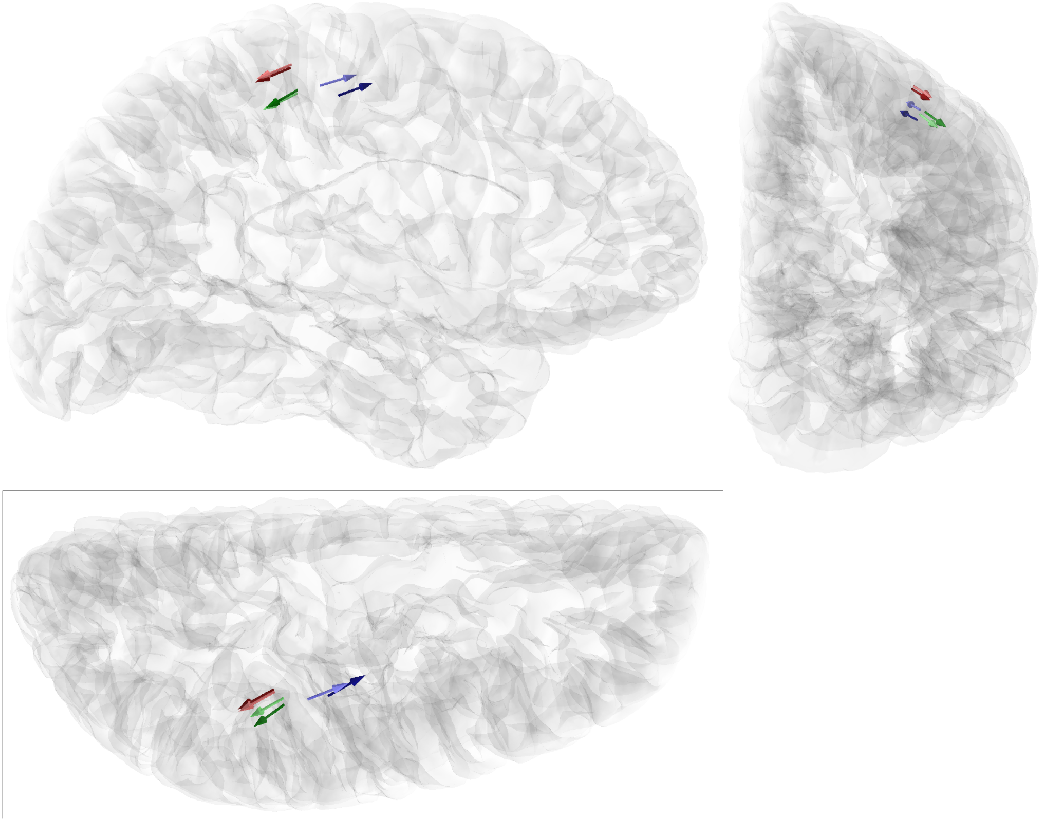
Locations of equivalent current dipoles representing the somatosensory responses N20m (blue), P35m (red), P60m (green) visualized on orthogonal views (lateral, caudal, dorsal) of the right hemisphere. Both OPM (light) and SQUID (dark)-based dipoles are shown.

Constraining the area in the MRI and optical surface scan used for co-registration is crucial to avoid artifacts. In addition, certain head surface features, such as the jaw, may be slightly different in the MRI and surface scan due to the difference in body orientation. Another evident difference is the compression of some facial features, especially in the cheeks, due to the padding used to minimize head movement during MRI acquisition. Facial areas such as the brow and forehead should function as more dependable areas for co-registration, as there the skin does not move as much in relation to the skull as in other areas.

Co-registration is not the only source of errors in source estimation. In our experiment with the phantom, inaccurate sensor calibration and the small number of channels may have hampered dipole localization. The phantom dipoles were localized, on average, with better than 2.5-mm accuracy, which is similar to what has been reported earlier when applying optical co-registration to whole-head SQUID-based MEG systems [14], [15] and significantly more accurate than these studies reported for electromagnetic digitizer-based co-registration.

Our somatosensory measurement revealed the typical sequence of evoked responses for such stimuli. The first response appeared at approximately 20 ms after stimulus onset (N20m). Both OPM- and SQUID-derived N20m dipole models localized in the anterior the wall of the postcentral gyrus, corresponding to the primary somatosensory cortex. The SQUID-based N20m dipole was located deeper along the gyral wall and more laterally than the OPM dipole (difference 5.0 mm). However, both dipoles were located in Brodmann area 3b. The sources of both P35m and P60m responses localized in the posterior wall of the postcentral gyrus, corresponding to Brodmann area 2. For these responses, the difference in dipole position between the OPM and SQUID data was very small (0.9 and 1.6 mm, respectively). The SQUID-localized responses are not necessarily the perfect ground truth: the co-registration process used for the OPM system should be more accurate than that of the SQUID system. Additionally, the OPM measurement may reveal additional neural activity not seen in conventional SQUID-MEG due to the large source-sensor distance, which limits spatial resolution.

Previous work has shown that intersession variability of P35m localization using SQUID-MEG can be ~5 mm [30], and N20m localization between SQUID-based MEG systems can have a variability of ~8 mm [31]. However, as argued by Solomon and colleagues ([30]), some of the variability in these results may stem from co-registration errors, as electromagnetic digitizer-based co-registration was used. Keeping these results in mind, the differences between OPM- and SQUID-derived dipole localizations in this work are small.

As the structured-light scanner acquires data at 30 Hz, it could be applied for continuous head tracking during measurements in the future. This could be done either using retroreflective markers attached to the head of the subject or using the entire digitized surface. Retroreflective markers would provide easily recognizable reference points that can be used to track head movements without any manual preprocessing. However, as the number of markers would be limited, they would have to be attached robustly to locations that do not move in relation to the head.

The methodology presented in this work can easily be applied to conventional SQUID-based MEG, and should have significant accuracy and speed benefits over the traditional digitizer-based approach. Similar methods applied to SQUID-based MEG have previously been presented by Bardouille and colleagues [14] and Murthy and colleagues [15].

In the future, as the number of sensors in OPM-based MEG system increases, optical surface scanning can also be used in conjunction with HPI coils for tried-and-true real-time head tracking. However, current-generation OPMs have a limited bandwidth only up to 150 Hz, meaning that only a small frequency window, which is close to that of physiological MEG data, is available for electromagnetic co-registration methods during MEG measurements.

## VI. Conclusions

We present a optical co-registration method that can be applied to on-scalp MEG as well as to current conventional MEG systems. The optical co-registration method is more accurate and faster than conventional electromagnetic digitizer-based methods.

## Acknowledgment

The authors thank the Advanced Magnetic Imaging centre of Aalto Neuroimaging Infrastructure for the MR images and Aalto University Science-IT for providing computational resources.

